# Confinement with Myosin-II suppression increases heritable loss of chromosomes, using live-cell ChReporters

**DOI:** 10.1101/2023.02.25.530049

**Authors:** Brandon H Hayes, Peter Kuangzheng Zhu, Mai Wang, Charlotte R Pfeifer, Yuntao Xia, Steven Phan, Jason C Andrechak, Junhong Du, Michael P Tobin, Alisya Anlas, Lawrence Dooling, Manasvita Vashisth, Jerome Irianto, Michael A. Lampson, Dennis E Discher

**Affiliations:** Mol. Cell Biophysics Lab, University of Pennsylvania, Philadelphia, PA 19104; Biology, University of Pennsylvania, Philadelphia, PA 19104

**Keywords:** Myosin, Rigidity, Copy Number, Heritability

## Abstract

Matrix around cells exerts many effects, some of which depend on the putative tumor suppressor Myosin-II, but whether such factors affect DNA sequences in a cell remains unclear. Here, live-cell monitoring of changes to chromosome copy numbers is developed and studied under diverse perturbations, including Myosin-II inhibition in confined mitosis. Squeezing of mitotic cells is seen *in vivo* and kills *in vitro*, but stem cells and cancer cells that survive show heritable loss of mono-allelic GFP/RFP-tagged constitutive genes that function as novel Chromosome-reporters (ChReporters). Myosin-II suppression increases such loss in 3D & 2D confinement but not in standard 2D, with “lethal” multipolar divisions proving myosin-dependent. Viable chromosome loss after confined mitosis associates more with mis-segregation than with multipolars or division number. Solid human tumors and teratomas in mice also show ChReporter loss and a confinement-signature of Myosin-II suppression, although losses are selected against in 2D culture. Heritable loss in rigid-confinement also appears independent of a spindle assembly checkpoint that functions in 2D. Confinement and myosin-II thus regulate pathways of heritable mechanogenetic change.

## INTRODUCTION

Mechanical aspects of a cell’s microenvironment such as matrix stiffness, confinement, and stretching can affect gene expression as well as the structure and function of cells in interphase [1-8]. Myosin-II is often key [9], but myosin-II also has roles in cell division such as mitotic rounding of animal cells that works against 3D compression (**Fig.1A**) [10, 11]. Intriguingly, deletion of myosin-II in rigid yeast somehow causes chromosome losses and gains that are heritable [12], and mouse embryo knockdown of non-muscle myosin-IIA in skin – which is a dense and stiff 3D tissue – reproducibly causes cancer [13, 14]. Because genetic changes typically drive cancer and are far more prominent in tumors that arise in solid tissues [15, 16], we hypothesized that myosin-II suppression can directly increase genetic changes – particularly when division is confined or constrained.

**Fig.1.**
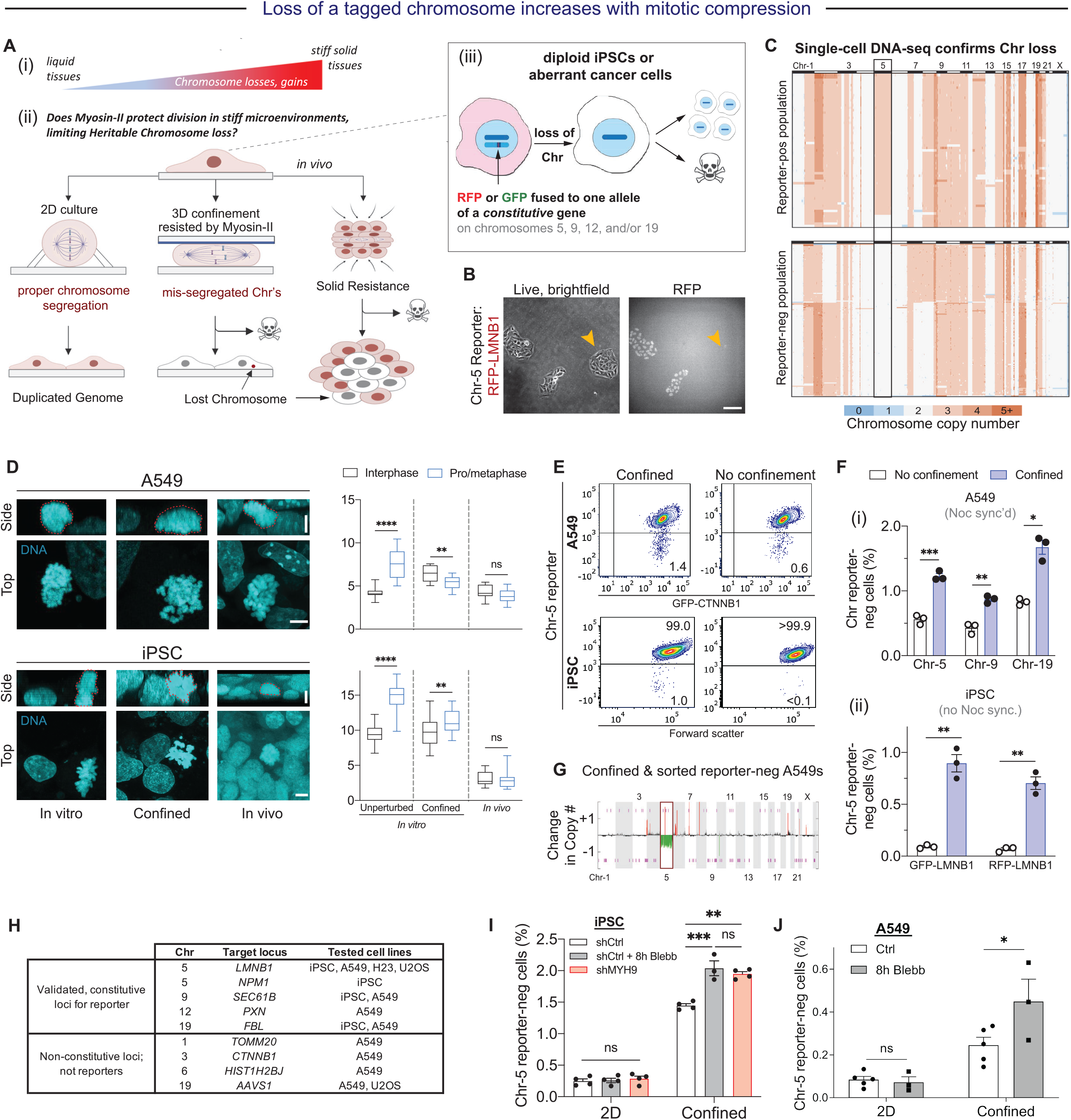
Live-cell tracking of monoallelic RFP/GFP fusions to constitutive alleles shows rigid mitotic confinement leads to chromosome loss. **(A)** Chromosome losses and other genetic changes are highest in tissues that are both stiff and proliferative **(i)**, motivating development of a live cell chromosome reporter (ChReporter). **(ii)** Mitotic perturbations potentially affect Chr loss and/or viability: maximal rounding in 2D culture is suppressed when cells are rigidly confined *in vitro* or surrounded *in vivo* by cells and matrix. **(iii)** Alleles are identified that are constitutively expressed even when fused to GFP or RFP. **(B)** RFP-positive and RFP-negative A549 colonies using the Chr-5 reporter in which one allele of *LMNB1* has an N-terminal RFP. A549 cells were sorted to purity via FACS, plated sparsely, and allowed to grow for a week. Scale bar = 100 µm. **(C)** Single-cell DNA sequencing reveals both expected and unexpected chromosome (Chr) losses and gains in A549 cells with the RFP-LMNB1 reporter, after isolation via FACS. Each row shows the whole genome at 1.5 Mb resolution for one of 61 RFP-pos or 140 RFP-neg cells. **(D)** Images of chromatin and plots of chromatin height in iPSCs and A549 cells in either standard 2D culture, rigidly confined culture, or 3D *in vivo* teratomas or tumors engrafted at subcutaneous sites in immunodeficient mice (n≥30 cells per condition, Mean and SEM; Mann-Whitney U-rank test: \*\**p* < 0.005; \*\*\*\**p* < 0.0001). Scalebars = 5 µm. **(E,F,G)** Flow cytometry analyses of Chr-5 reporter loss in iPSCs and A549s in either confined or standard 2D cultures: **(i)** Noc-synchronized A549 cells with three different reporters, or **(ii)** two distinct iPSC clones with no synchronization because iPSCs double in ∼10h, which is ∼3x faster than A549s. (n = 3 replicates, Mean and SEM; unpaired two-tailed t-test with Welch’s correction: \**p* < 0.05, \*\**p* < 0.005, \*\*\**p* < 0.0005). Flow sorted cells are isolated immediately after confinement to validate ChReporter loss in A549 cells based on SNPa analyses. **(H)** Designs for various loci and cell lines, with some proving to be constitutively expressed and validated ChReporters. **(I,J)** Flow cytometry results for ChReporter-negative iPSCs and imaging results for A549s, which both show increases under rigid confinement with myosin-IIA suppression (knockdown or inhibition with blebbistatin) but no effect in standard 2D. Mean and SEM (n = 3 or 4; 3-way ANOVA with Tukey’s correction for multiple comparisons: \**p* < 0.05, \*\**p* < 0.005; \*\*\**p* < 0.0005).

Division under compression distorts the spindle of microtubules (MT) and acutely mis-segregates chromosomes but also kills some cells quickly [17-19]. Heritable losses or gains in chromosomes (i.e. aneuploidy) might eventually emerge, but viable changes remain unexamined and nontrivial given the cell death in mitotic confinement that includes striking “*lethal multipolar divisions*” [18]. Constraints imposed by 2D micropatterns helped show that myosin-II inhibition with blebbistatin perturbs centriole separation, leading the authors to speculate that myosin-II “*might act in fundamental mechanisms of aneuploidy prevention*” [20]. Although blebbistatin in initial studies of standard 2D cultures “inhibited contraction of the cleavage furrow without disrupting mitosis” [21], subsequent studies with this reversible inhibitor and with knockdowns concluded that division is perturbed [22-24], but only one recent study claimed increased frequency of abnormal chromosome segregation, suggesting “*An interesting possibility is that a lack of the acto-myosin network is correlated with aneuploidy*” [25]. Thus, roles for myosin-II in heritable aneuploidy – coupled or not to 2D or 3D microenvironments – remain speculative but perhaps important to oncogenesis.

To visualize heritable loss of a chromosome (and potentially a gain) by a novel approach that allows verified tracking of viability (*rather than death*), a candidate ‘constitutive’ gene on a chromosome (i.e. Chr-5, 9, 12, &/or 19) is gene-edited as a GFP/RFP fusion to create a ‘ChReporter’ (**Fig.1A**). Loss of the GFP/RFP signal allows us to then identify individual cells and colonies *before, during, and after* the change – with Chr loss for some tagged genes (not all) made clear by end-stage genetic analyses. Viability and heritability are thus definitive rather than inferred from sequencing of DNA/RNA extracted from dead cells. Our approach is compatible with many studies in cell biology, and overcomes error rates in sequencing (e.g. [26]) that add uncertainty in rare cell detection (e.g. ∼0.1 to 1% of cells). High sensitivity is crucial to understanding initial changes that give rise via eventual selection to new cell populations. For studies of confinement-induced ChReporter loss, we focus on solid tumor derived lines (e.g. A549 lung adenocarcinomas) and a line of genomically-stable induced pluripotent stem cells (iPSCs) as a model stem cell relevant to normal tissue and tumor stem cells [27, 28]. Heritable changes are ultimately shown to be a genuinely genetic mechano-response to constraining microenvironments.

## RESULTS

### Myosin-II mechanoprotects against ChReporter loss in confined mitosis

Within days of a fresh flow cytometry sort for cells with a specifically targeted organelle such as the nucleus with RFP-LMNB1, loss of GFP/RFP is evident in very rare cells and colonies (∼0.1%) (**Fig.1B**). Such colonies provide evidence of viability and heritability, and sorting then allows assessment of the molecular basis of ChReporter loss by single-cell sequencing, bulk array methods, and low-throughput karyotype spreads among other approaches (**Fig.1C**). Detecting loss of a given gene (e.g. GFP/RFP) at high accuracy as required for our studies remains challenging, but sorted A549s subjected to single-cell sequencing not only shows the expected ChReporter loss but also other unexpected losses and gains in rare cells. This evidence of spontaneous but low genetic instability in A549s contrasts with the relative lack of ChReporter-neg iPSCs in standard 2D culture, which again agrees with single-cell sequencing that shows the line is genomically-stable until perturbed.

To first address the physiological relevance of mitotic compression, we analyzed confocal stacks within human-in-mouse tumors and teratomas. Both mitotic chromatin and interphase nuclei show the same height in iPSC-derived teratomas and A549 tumors -- unlike standard 2D cultures (**Fig.1D**). The teratomas have a rigidity that is palpably similar to the tumors, which are collagen-rich and stiff (∼5 kPa) [29] and also within a stiffness range associated with high genetic change in tumors [30, 31]. Rigid confinement of cultures was therefore used to suppress the typical mitotic rounding in 2D cultures and thereby mimic stiff tissue: a ring-weight was placed on top of an upper glass coverslip with compression limits provided by rigid polystyrene microbeads mixed with the cells. Confinement even for ∼8h suffices to visibly increase abnormal mitosis, with mis-segregated chromosomes and suppressed growth (**Fig.1D**) as well as cell death indicating intolerable stress (**Fig.1A**).

Viable RFP/GFP-negative cells were quantified by flow cytometry after 16-48h of recovery in 2D culture following confinement. Approximately 1% of A549s and of iPSCs show loss of various ChReporters -- which is ∼2-10 fold higher than 2D culture controls (**Fig.1E,F**). Flow sorting of RFP-neg A549s engineered with a Chr-5 ChReporter was followed by genetic analyses that show the expected Chr loss relative to RFP-pos controls (**Fig.1G**). Loss of diverse ChReporters was also validated for multiple cell types, but an important aspect of the method is that some tagged genes clearly show GFP/RFP-neg cells *do not represent* a genetic change (**Fig.1H**).

To test whether myosin-II might protect against ChReporter loss, we first knocked down the major non-muscle myosin-II isoform, myosin-IIA *(MYH9*), in rapidly dividing iPSCs that harbor the Chr-5 ChReporter. The 3D-compressed knockdown cells showed 30-40% more ChReporter loss than control cells in confinement but with no effect in 2D (**Fig.1I**). To rule out adaptation to the knockdown and to inhibit other non-muscle myosin-II isoforms in these cells, we added the pan-myosin-II inhibitor blebbistatin only during the 8h confinement; such a brief treatment affects the levels of very few proteins compared to knockdown [32]. Blebbistatin treatment nonetheless showed the same quantitative effect as knockdown: no effect in 2D and significant increase under 3D-compression. A549 cells confirmed the result (**Fig.1J**).

Given that sh*Myh9* knockdown in embryonic mouse skin by Fuchs & coworkers led to carcinoma [13] that first suggested a tumor suppressor role for myosin-II [33], and also given that skin is a relatively stiff and constraining 3D microenvironment [30, 31], the results here provide the first evidence of genetic changes (per cancer) in myosin-II suppressed cells and *only* in confinement. Such changes have remained unclear in the field as has the relevance to human cells

### Confinement induced mis-segregation is skewed by myosin-II inhibition

Blebbistatin has no effect on a confinement-induced suppression of anaphase counts (**Fig.2A**), with past studies also showing confinement-induced delays in progression to anaphase [17, 18]. Tubulin staining confirms a significant confinement-induced lengthening of spindle [17, 18], and we observe blebbistatin does not affect the results. Interphase cells have a more dendritic shape with blebbistatin, which is typical, but blebbistatin had no effect on the shape or overall F-actin signal from rhodamine-phalloidin in mitotic cells; a difference between actin and myosin-II is suggested because latrunculin in confinement strongly *increases* “lethal” multipolars as well as cell death [18]. Contrary to the one report that blebbistatin increases chromosome mis-segregation in 2D culture [25], we observe no effect of myosin perturbations in 2D culture on bridging chromosome events (**Fig.2B,C**). In 3D confinement, however, the large fraction of lethal multipolar events [18] is rescued by blebbistatin (**Fig.2D**), consistent mechanistically with blebbistatin’s inhibition of centriole separation [20]. Abnormal anaphase counts are overall unaffected by blebbistatin (**Fig.2C**), and so blebbistatin in 3D confinement increases the more tolerated bridging events, with no other obvious effects of blebbistatin other than the suppression of the lethal multipolars. Thus, the net increase in tolerable mis-segregation events can explain the ChReporter loss with myosin-II inhibition in confinement (**Fig.1J**).

**Fig.2.**
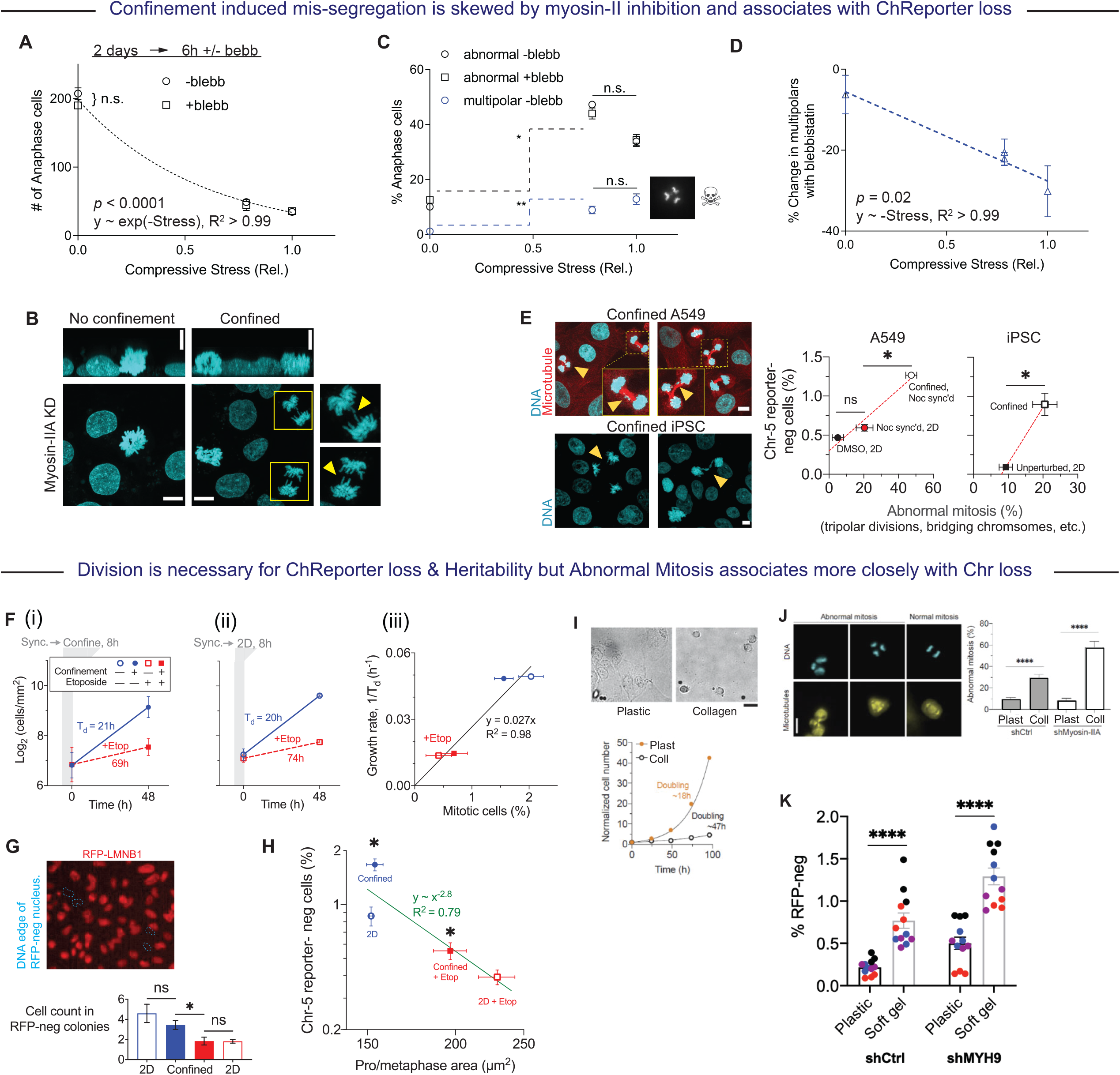
Confined division with myosin-II inhibition increases viable Chr mis-segregation and ChReporter loss. **(A)** A549 cells plated for 2 days were compressed per Fig.1 or with slightly lower stress for 6 h ± blebbistatin and then fixed, stained, and imaged for anaphase counts. Blebbistatin has no effect on the effect of mitotic compression. (n = 3-6 replicates, Mean and SEM; with log-linear model for non-zero slope *p* < 0.0001, with exponential fit R_2_>0.99). **(B)** MYH9 knockdown shows abnormal mitoses under rigid confinement (yellow arrowheads for bridging chromosomes), whereas unconfined iPS cells only show normal mitoses, indicating that shMYH9 does not affect basal instability. Scale bar = 10 µm. **(C,D)** Abnormal anaphase increases with compression but is always independent of blebbistatin (n = 3-6 replicates, Mean and SEM; unpaired two-tailed t-test with Welch’s correction: \**p* < 0.05, \*\**p* < 0.005). Multipolars are included as abnormal and are suppressed by blebbistatin. (n = 3-6 replicates, Mean and SEM; for non-zero slope *p* = 0.02, with linear fit R_2_>0.99). **(E)** Images of abnormal mitosis and flow cytometry measures of the percentage of Chr-5 reporter-neg cells plotted against percentage of abnormal mitosis for A549s and iPSCs. (n = 3 replicates, Mean and SEM; unpaired two-tailed t-test with Welch’s correction: \**p* < 0.05). Scalebar = 5 µm. **(F-H)** TOP2A inhibition with Etoposide during confinement of synchronized A549. Proliferation remains suppressed even after drug washout. Cell numbers in colony forming units (CFU) of ChReporter-neg cells, which Etop suppresses. (n = 3 replicates, mean and SEM; two-way ANOVA with Tukey’s correction for multiple comparisons: \**p* < 0.05). **(I)** A549 cells spread on plastic but are always round and laterally confined while attaching firmly on dense collagen-coated substrate. Scalebar = 50 µm. **(J)** Abnormal mitosis is more frequent on collagen-coated substrate versus plastic, in both Myosin-IIA depleted cells and shCtrl cells. Mean and SEM (n ≥ 10 cells per condition; unpaired two-tailed t-test with Welch’s correction: **** p<0.0001). Scale bar = 20 µm. **(K)** ChReporter RFP-neg cells occur more frequently on collagen-coated gels versus plastic. (n = 4 triplicate studies; 3-way ANOVA with Tukey’s correction for multiple comparisons: \*\**p* < 0.005; \*\*\**p* < 0.0005).

Mitotic chromatin is flattened and compacted by the compression (**Fig.2B**), whereas interphase cells are not as tall and are unaffected. ChReporter-neg cells generally associate with visible levels of abnormal mitosis (**Fig.2E**), although abnormal mitosis does not strictly predict Chr loss: some iPSCs show abnormal mitosis but no ChReporter loss whereas A549s show the opposite. ChReporter loss is far below the fraction of cells exhibiting abnormal mitosis (∼15-fold for iPSCs, and ∼60-fold for A549s), which seems consistent with losses of untagged chromosomes as well as Chr-9 and Chr-19 ChReporter results (**Fig.1F**). Some studies used transient nocodazole (Noc) to synchronize A549 division before confinement, and a residual effect (after drug washout) fits the trend (**Fig.1H**) in agreement with known perturbations by Noc-induced disassembly of spindle MTs [34].

To assess the role of cell division, the Topoisomerase-IIα inhibitor Etoposide (Etop) was added at a low non-toxic dose [35] during the 8 h confinement. Etop suppresses subsequent growth and mitotic counts (**Fig.2F**) as well as ChReporter-neg cells and colony size that indicates heritability (**Fig.2G,H**). Viable A549s re-spread their decondensed chromatin in interphase, re-assemble their lamina, and proliferate normally (without Etop) to generate ChReporter-neg colonies similar in size to 2D cultures (**Fig.2G**). Isolated chromosomes are physically softened by topoisomerase TOP2A [36], and in 2D cultures, TOP2A drives compaction of mitotic chromatin [37, 38], which thus remains spread in the presence of Etop (**Fig.2G**); however, compressed mitosis modestly rescues this compaction defect and also increases ChReporter loss. The latter effect is nonetheless small, and so the Etop results suggest that more divisions *N* tend to favor more chromosome loss.

2D substrates with a dense collagen-coating can limit cell spreading [39]. We thought that this might allow us to test the effects of lateral confinement on ChReporter loss, especially in light of recent results from 2D micropattern constraints that led to the speculation that myosin-II “*might act in fundamental mechanisms of aneuploidy prevention*” [20]. We first discovered that A549 proliferation is greatly slowed on collagen-coated substrates relative to standard 2D culture (i.e. fewer divisions *N*) (**Fig.2I**), and then we quantified a large increase in abnormal mitosis (**Fig.2J**), which aligns well with 3D confinement effects (**Fig.2A-C**). ChReporter loss relative to 2D cultures increases similarly (**Fig.2K**). Moreover, Myosin-IIA suppression tends to show more abnormal mitosis and more ChReporter loss relative to controls, consistent with a recent speculation about its role in aneuploidy [20]. Abnormal mitoses induced by confining microenvironments thus associate better with Chr loss than with divisions *N*. Solid tissues such as skin are also rich in collagen, which makes them stiff [29] and which leads to a prediction of Myosin-II suppressed ChReporter loss within stiff and constraining solid tissues *in vivo*.

### *In vivo* Chromosome loss increases with rigidity-associated divisions & Myosin-II suppression

To directly assess 3D *in vivo* responses of ChReporters including a conceivable role for Myosin-II, subcutaneous xenografts were made in stiff dermis, which is rich in mouse-collagen [29]. Immunodeficient mice were injected with human iPSCs or A549s expressing *LMNB1* Chr-5 ChReporters. iPSC teratomas and A549 tumors (**Fig.3A**) harvested after ∼2-3 months were flow-sorted for the rare fluorescent-negative cells that confirm Chr-5 loss by genetics analyses (**Fig.3B**). The result rules out rare epigenetic changes [40]. Disaggregated teratomas and tumors further confirm rare human cells lack nucleus-localized GFP/RFP-LMNB1 (**Fig.3C**). Importantly, flow cytometry showed ChReporter-negative cells increased in all teratomas and tumors (**Fig.3D-G,top**), ranging from 2- to 30-fold more loss of RFP/GFP-LMNB1 than time-matched or long-term 2D culture controls.

**Fig.3.**
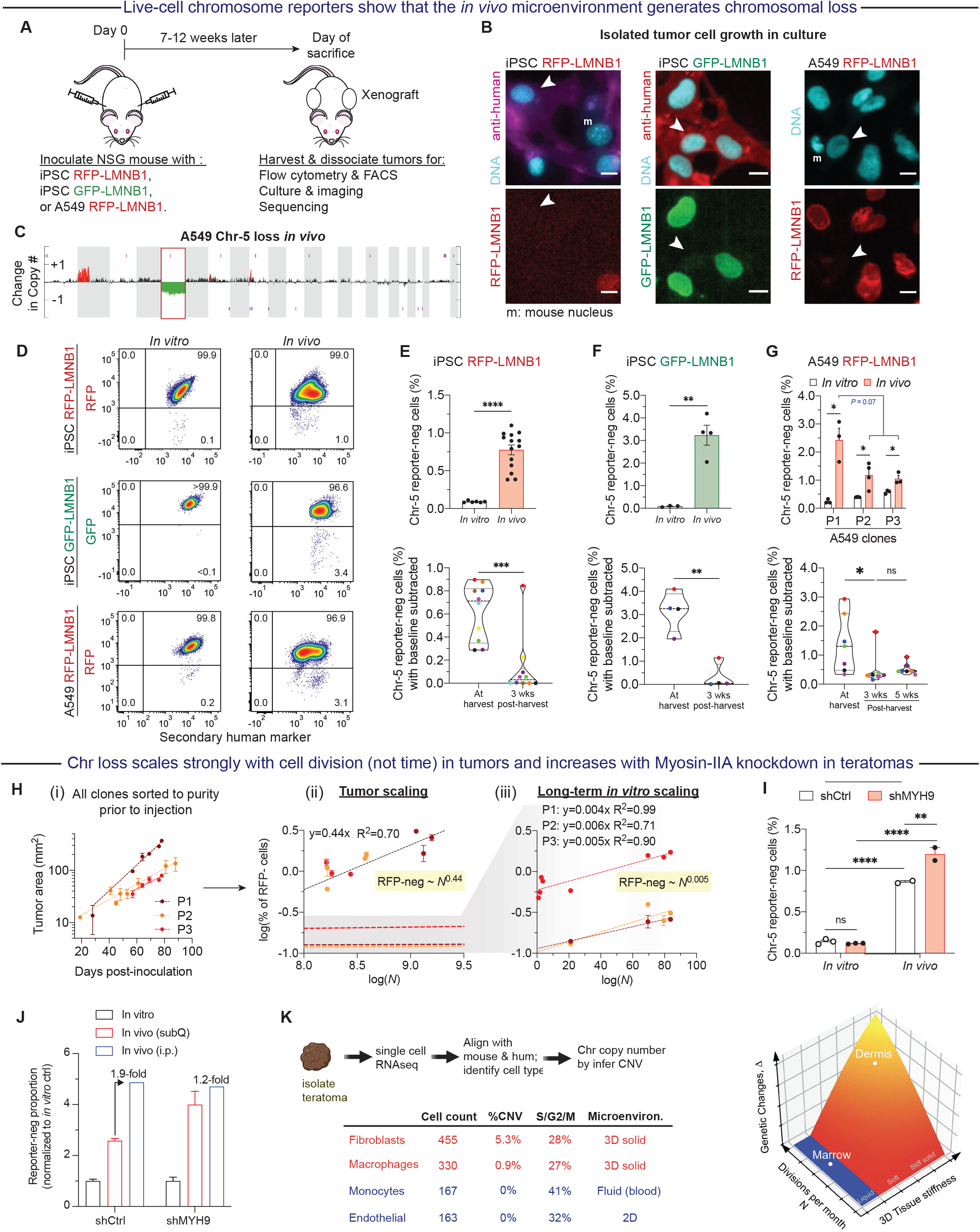
Chromosome losses increase in 3D microenvironments *in vivo* with modulation by Myosin-II. **(A-C)** Human cells xenografted at subcutaneous sites in immunodeficient NSG mice. When the iPSC teratomas or A549 tumors grew to a diameter of ∼2 cm, they were harvested, disaggregated, and analyzed for Chr-5 reporter loss. SNPa validation of FACS-sorted RFP-neg A549 cells confirm reporter loss indicates chromosome loss (n = 3 tumors). Images of Chr-5 reporter loss (LMNB1 protein) in 2D-cultures (∼1 wk) of cells derived from iPSC teratomas (both clones) or from A549 tumors. Mouse cells show distinctive chromocenters in Hoechst stain of DNA. Scale bar = 10 µm. **(D-G)** Flow cytometry shows increased Chr-5 reporter-negative loss (%) from *in vivo* harvested cells compared to time-matched, 2D culture controls for all teratomas and tumors. (H-K) Quantitation of Chr-5 reporter-negative cells from various teratomas or tumors versus *in vitro* cultures, including 3-5 wk cultures post-harvest for assessments of persistent viability. Mean and SEM (n = 3-14 replicates; unpaired two-tailed t-test with Welch’s correction between each confinement condition and its respective unconfined, standard 2D control: \*\**p* < 0.005, \*\*\**p* < 0.0005, \*\*\*\**p* < 0.0001; ns, not significant). **(H)** Tumor growth for all three A549 clones plotted vs time **(i)** or ChReporter-neg’s plotted vs cell division number *N* **(ii)**. The latter is estimated from tumor size and mean cell volume per confocal imaging of tumors (Fig.1I). Power-law scaling of Chr-5 reporter-negative A549 cells is much stronger *in vivo* than in 2D cultures **(iii)**, as measured over 200 days with weekly splitting. **(I)** Myosin-IIA knockdown iPSCs *in vivo* show the most Chr-5 reporter-negative cells (RFP-LMNB1) upon disaggregation of solid teratomas (of shMYH9) when compared to *in vivo* controls (shCtrl) or time-matched standard 2D-cultures (*in vitro*). Mean and SEM (n = 2-3; two-way ANOVA with Tukey’s correction for multiple comparisons: \*\**p* < 0.005; \*\*\*\**p* < 0.0001). **(J)** Fold-change in ChrReporter-negative cells generated in teratomas (with or without Myosin-IIA knockdown) versus corresponding passage-matched 2D controls. All reporter-loss percentages are normalized to the respective 2D control. Stiff teratomas at subcutaneous (subQ) sites show more ChReporter-neg’s with Myosin-IIA knockdown, unlike soft teratomas at intraperitoneal (IP) sites. Mean and SEM (n = 2-3; two-way ANOVA with Tukey’s correction for multiple comparisons: \*\**p* < 0.005; \*\*\*\**p* < 0.0001). **(K)** Normal, solid tissue mouse cells exhibit rare and shared chromosome loss or gain based on single-cell RNA-sequencing of teratomas Freshly isolated teratoma was disaggregated and split for single cell RNA-seq, with species determined by alignment to reference genomes for human (GRCh38) or mouse (GRCm38). Gene expression profiles were then used for cell type annotation using singleR and copy number from inferCNV (Tickle et al., 2019). (**Plot**) Genetic changes increase with divisions and with 3D rigidity of the microenvironment.

Heritability is again limited after stress removal. ChReporter-neg cells *ex vivo* are generally out-proliferated by ChReporter-pos cells across multiple 2D cultures. Teratoma-derived ChReporter-neg cells mostly died by 3 weeks, with crucial exceptions of viable cells from two teratomas (**Fig.3E,F-bot**); infrequent genetic changes in iPSCs limit their use [27]. ChReporter-neg A549s from tumors also decreased in frequency by 3 wks of culture but then tended to grow (**Fig.3G-bot**), consistent with robust persistence of abnormal cancer cells. Genetic change under the distinct stresses of 3D is nonetheless highlighted by the uniformly higher percentages of ChReporter-neg cells from freshly harvested teratoma/tumor cells versus 2D cultures of the same cells. Proliferation under 3D stress *in vivo* is a likely determinant because differences in the percentage of RFP-neg cells between the three A549 RFP-LMNB1 clones (**Fig.3G-top**) correlate with the distinct growth rates of the tumors (**Fig.3H-i**). Indeed, cell volume estimates from confocal images (**Fig.1D**) allow us to convert measured tumor sizes at harvest to total cell cycle numbers (*N*), yielding a much stronger power-law of [%RFP-neg] ∼ *N*^a^ (*a* = 0.44) for *in vivo* relative to standard cultures where cells round up and divide unstressed by the overlying fluid and [%RFP-neg] ∼ *N*^*b*^ (*b* = 0.005) (**Fig.3H-ii,iii**).

Myosin-IIA knockdown of *LMNB1*-edited iPSCs showed ∼50% more loss in teratomas than controls (**Fig.3I**), consistent with *in vitro* effects in rigid confinement of iPSCs (**Fig.1I**). Solid teratoma masses have the same consistency as subcutaneous tumors rich in mouse-derived collagen and mouse cells [29]. However, intraperitoneal xenografts lack solidity, and intraperitoneal teratomas show no increase in ChReporter loss upon myosin-IIA knockdown (**Fig.3J**). The findings offer insight into how a similar myosin-IIA knockdown in the stiff dermis of mouse embryo leads to cancer [13].

To determine whether any normal mouse cells in the teratomas also exhibit Chr copy number variations (CNVs), we applied single cell RNA-seq, using sequence differences to identify various mouse lineages (fibroblasts, endothelial cells, and immune cells; see Methods for inferCNV). Importantly, tissue micro-environments for fibroblasts and macrophages are 3D and matrix-rich and thus likely to confine mitosis, and both lineages show ∼1-5% of cells with CNVs (**Fig.3K**). In contrast, the other two detected lineages show no CNVs despite evidence of replication: endothelial cells proliferate in 2D monolayers and thus mitotically round up with an overlying fluid that should maintain the fidelity of Chr segregation. Past work showing genetic changes in normal tissue has been unclear in terms of lineage specificity and such 2D versus 3D microenvironments. Results here thus suggest genetic changes Δ for replicating cells are much lower in soft or fluid tissues that include 2D layers (e.g. endothelia) versus cells in stiff 3D tissues.

### Inhibiting spindle assembly checkpoint increases ChReporter loss in 2D but not 3D confinement

To assess whether other factors or pathways affect outcomes of 3D-rigid confinement, we studied inhibitors of the spindle assembly checkpoint (SAC) that control exit from pro-metaphase in 2D [41-44]. Chr losses and gains in 2D cultures are well known to be induced by inhibiting MPS1 kinase in the SAC, but unlike confined mitosis that distends the spindle [17] and spreads mitotic chromatin, MPS1 inhibition (MPS1i) has no effect on mitotic spreading in 2D culture of A549s. In 2D, MPS1i causes abnormal mitosis as expected [43, 44] and also causes dose-dependent ChReporter loss (**Fig.4A-i**). In confinement, MPS1i had surprisingly no effect on ChReporter loss in A549s (**Fig.4A-ii**), with abnormal mitosis remaining *equally* high in confinement regardless of MPS1i (**Fig.4A-iii**). A second drug confirms the results (AZ3146: **Fig.4B**). Despite different outcomes in 2D versus 3D, the prolonged mitosis in confinement relative to 2D controls [17-19] is accelerated by MPS1i to the same extent in 2D and 3D (**Fig.4A-iii**). This is consistent with sustained knockdown of the SAC component, MAD2 [18]. The main difference with mitotic confinement is that cell death (often in pro-metaphase) is a major alternative fate even without drugs. While cell death is a major fate with some drugs that affect mitosis in 2D [45], the stress of confinement can be lethal in combination: MPS1i kills all confined iPSCs. Also, MPS1i in 2D consistently yields ChReporter loss of ∼1% that is similar to 3D-rigid confinement for diverse ChReporters in various cell types, and high doses cause cell death (**Fig.4C**).

**Fig.4.**
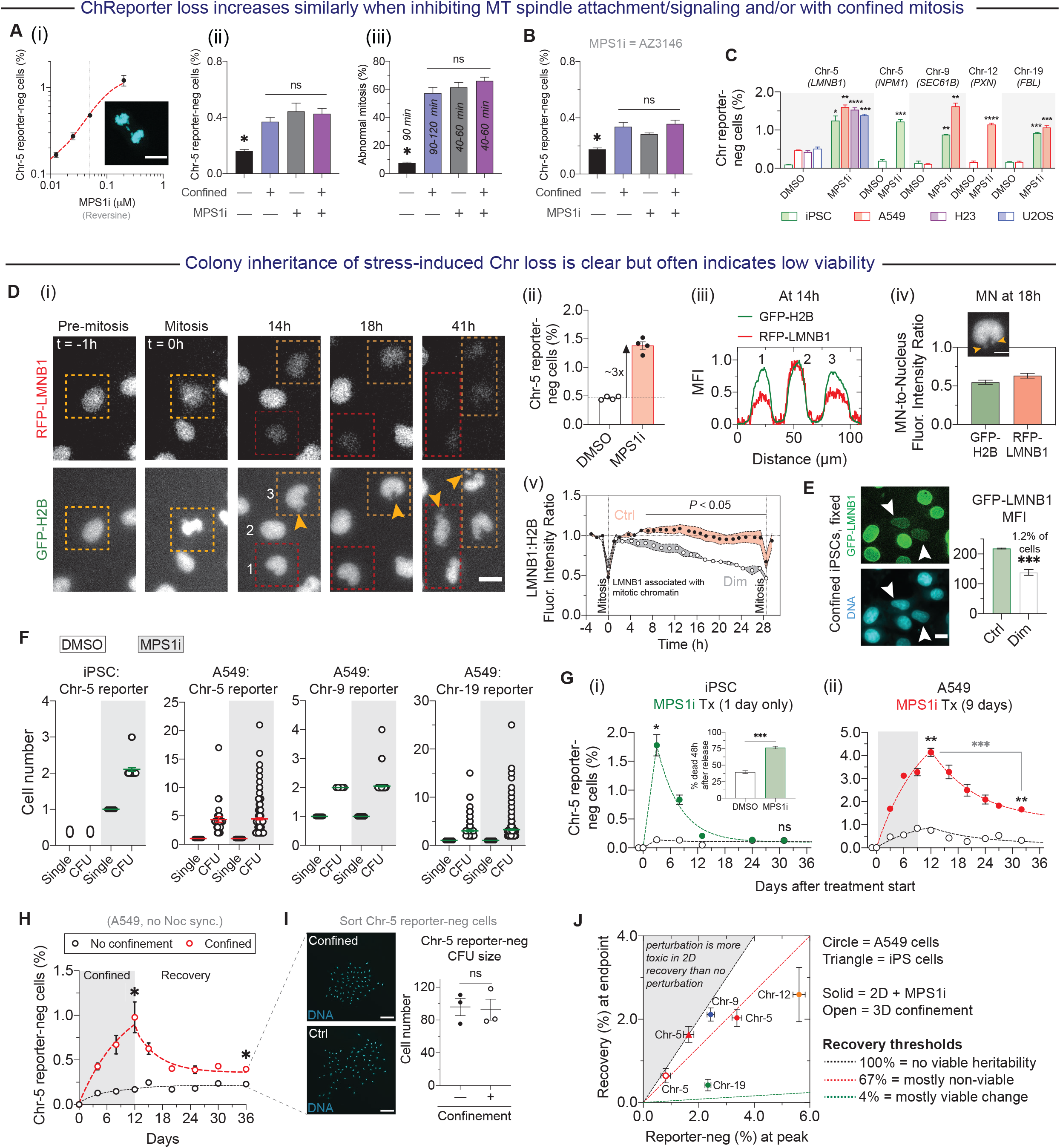
Confinement-induced ChReporter loss is unaffected by inhibition of a ‘2D’ mitotic checkpoint, and Colony inheritance of Chr loss often indicates relatively low viability during 2D recovery. **(A)** Confined mitosis has the same effect on ChReporter loss in A549 cells as inhibition of microtubule spindle attachment/signaling with MPS1 inhibitor reversine. **(i)** Saturable dose response with MPS1i (n = 3 replicates, Mean and SEM). Scalebar = 10 µm. **(ii)** Sub-saturating MPS1i, confinement, or a combination for 8h per day over 4 days show the same ChReporter effects, and the same (**iii**) Abnormal Mitosis%. Time ranges for mitosis indicate MPS1i rescues the prolonged division caused by confinement. (n = 3 replicate, mean and SEM, for both (ii) and (iii); two-way ANOVA with Tukey’s correction for multiple comparisons: \**p* < 0.05). **(B)** Second MPS1 drug AZ3146 used at a dose that gives similar ChReporter loss as confinement confirms the findings of A-ii. (n = 3 replicates, mean and SEM; two-way ANOVA with Tukey’s correction for multiple comparisons: \**p* < 0.05). **(C)** ChReporter-neg cells for all engineered lines (iPSCs, A549 and H23 lung adenocarcinoma, and U2OS osteosarcoma) treated with MPS1i or DMSO for 3 days. (n = 3 replicates, mean and SEM; unpaired two-tailed t-test with Welch’s correction: \**p* < 0.05, \*\**p* < 0.005, \*\*\**p* < 0.0005). **(D)** Low-light imaging over 48h shows **(i)** A549s lose RFP-LMNB1 (expressed from Chr-5) but not GFP-H2B (expressed from Chr-6). Cells treated with low-dose reversine were treated for 24h and were then imaged every 20 min. Boxed cells show dimming RFP signal as they divide twice. Yellow arrowheads: micronuclei. Scalebar = 20 µm. **(ii)** Imaging quantification of Chr-5 reporter-neg A549 cells, with a minimum of 2,000 cells per well counted. **(iii)** Intensity profiles across three cells: cells 1 and 3 are RFP-dimming cells, while cell 2 remains RFP-positive. **(iv)** Micronuclei intensities for both RFP-LMNB1 and GFP-H2B. Scalebar = 10 µm. **(v)** Fluorescence intensity ratio between RFP-LMNB1 and GFP-H2B for RFP-positive control cells and RFP-dimming cells, with intensities normalized to pre-mitosis images. Cell cycles were adjusted to the mean cell cycle time (28.5h). Mean and SEM (unpaired two-tailed t-test with Welch’s correction between control and dimming conditions at the same time point: \**p* < 0.05). **(E)** Imaging of iPSCs (with GFP-tag on *LMNB1*) to identify reporter loss while undergoing mechanical confinement. DNA stain is not different between dimming and control cells, but the extent of GFP dimming is consistent with the time average in panel (D). Mean and SEM (unpaired two-tailed t-test with Welch’s correction between control and dimming conditions at the same time point: \*\*\**p* < 0.0005). Scalebar = 10 µm. **(F)** Cell numbers in reporter-negative colony forming units (CFUs) of iPSCs (with Chr-5 reporter) or A549s (with Chr-5, Chr-9, or Chr-19 reporters) when treated with a continuous low-dose reversine or control for 3-5 days. **(G,H,I)** Heritable Loss Model fits of Chr-5 reporter-negative kinetics in iPSCs or A549s after MPSi treatment and recovery, or for A549s after repeated cycles of rigid confinement for 12 days and recovery in 2D culture. For the latter after 36 days, flow-sorted RFP-neg cells were plated back sparsely 1:1 with RFP-pos cells; RFP-neg cell numbers in CFUs after 1 week are the same, and the mixture also showed the same total cell numbers for all RFP-pos and -neg sample conditions. Mean and SEM (n = 3 replicates; unpaired two-tailed t-test with Welch’s correction between each treatment condition at the same timepoint: \**p* < 0.05, \*\**p* < 0.005). Scalebar = 100 µm. **(J)** Heritability of peak loss of reporters and extent of decay from that peak during recovery. Data is from indicated reporters with control conditions subtracted.

To further assess perturbations of confined mitosis, we took advantage of findings in 2D cultures that show mitotic cells arrest when cyclin-B degradation is inhibited [46]. In 3D rigid confinement we find such inhibition has surprisingly no effect on the protracted mitotic exit, but it does suppress anaphase and ChReporter loss. We tentatively conclude that confinement-induced chromosome loss requires mitotic exit via anaphase.

Live imaging over days to track *de novo* ChReporter loss (**Fig.4D**) provides direct evidence of heritability in viable colonies, while also ruling out selective growth of pre-existing rare cells. This is critical because in development, Chr gains and losses that emerge do not survive [30]. Image analysis of >1k cells also confirms the ∼3-fold increase in RFP-LMNB1 loss with MPS1i (per **Fig.4A**; using low illumination to minimize photobleaching and DNA damage). Nuclear envelope breakdown in mitosis disperses RFP-LMNB1, and RFP then dilutes through progressive divisions (**Fig.4D**). In iPSCs after confinement, GFP-LMNB1 shows a similar dimming in ∼1% of cells (**Fig.4E**), but cells per colony forming unit (CFU) is much lower for iPSCs than A549s after the same interval (**Fig.4F**), even though iPSCs normally divide faster.

Imaging shows micronuclei (**Fig.4D, arrows**) that are a signature of mis-segregated Chr’s and that often accumulate DNA damage, especially when lamin-B is low [47, 48] and curvature is high [49]. A549s show more micronuclei post-confinement than iPSCs. This could reflect near-zero tolerance of iPSCs to Chr loss in 2D-culture without MPS1i (**Fig.1F,I, 3E,F**), and the many MPS1i-induced Chr losses or gains at short times disappear after withdrawing drug (**Fig.4G-i**).

### Cycles of confined mitosis show losses scale, but death limits heritability

Various ChReporter A549s in 2D show basal levels of ChReporter-negs and colonies even after 2 wks, and 9 days of MPS1i not only tends to increase ChReporter-negs and colonies but drug withdrawal also leads to variable levels of viable cells with sustained loss of all ChReporters (**Fig.4G-ii**). Importantly, compression-generated ChReporter-neg A549s are similar: once 3D-confinement cycles are stopped by switching to standard 2D culture, ChReporter-neg’s diminish to a level above controls (**Fig.4H**). Further culturing shows RFP-neg cells from all conditions exhibit heritable loss of ChReporter evident in equally large, viable colonies (**Fig. 4I**). The results are not only consistent with *de novo* genetic change but also with similar outcomes for MPS1 inhibition and mitotic compression.

Mathematically, a *Heritable Loss Model* (HLM) accounts for ChReporter loss rate and proliferation (**Fig.5A**), indicating slower net proliferation after loss (by 7% to 47%). Rigid 3D-confinement and MPS1i accelerate Chr-5 loss by ∼4-fold relative to controls. The HLM also accommodates power laws for ChReporter loss versus divisions *N* (**Fig.5B**), as applied *in vivo* (**Fig.3H**). For 2D cultures where stiffness *E* = 0 for the overlying fluid phase, ChReporter loss Δ ∼ *N*^*a*^ gives *a* ∼ 0.03 to 0.1 (i.e. weak scaling). For rigid confinement (*E* >> 0) and for MPS1i perturbations, Δ ∼ *N*^*b*^ with *b/a* ∼ 2.5. The modeling consistently shows stress-driven acceleration of Chr loss (by ∼2 to 10-fold), but then growth or viability in 2D also shows Chr- and cell-type specific differences.

**Fig.5.**
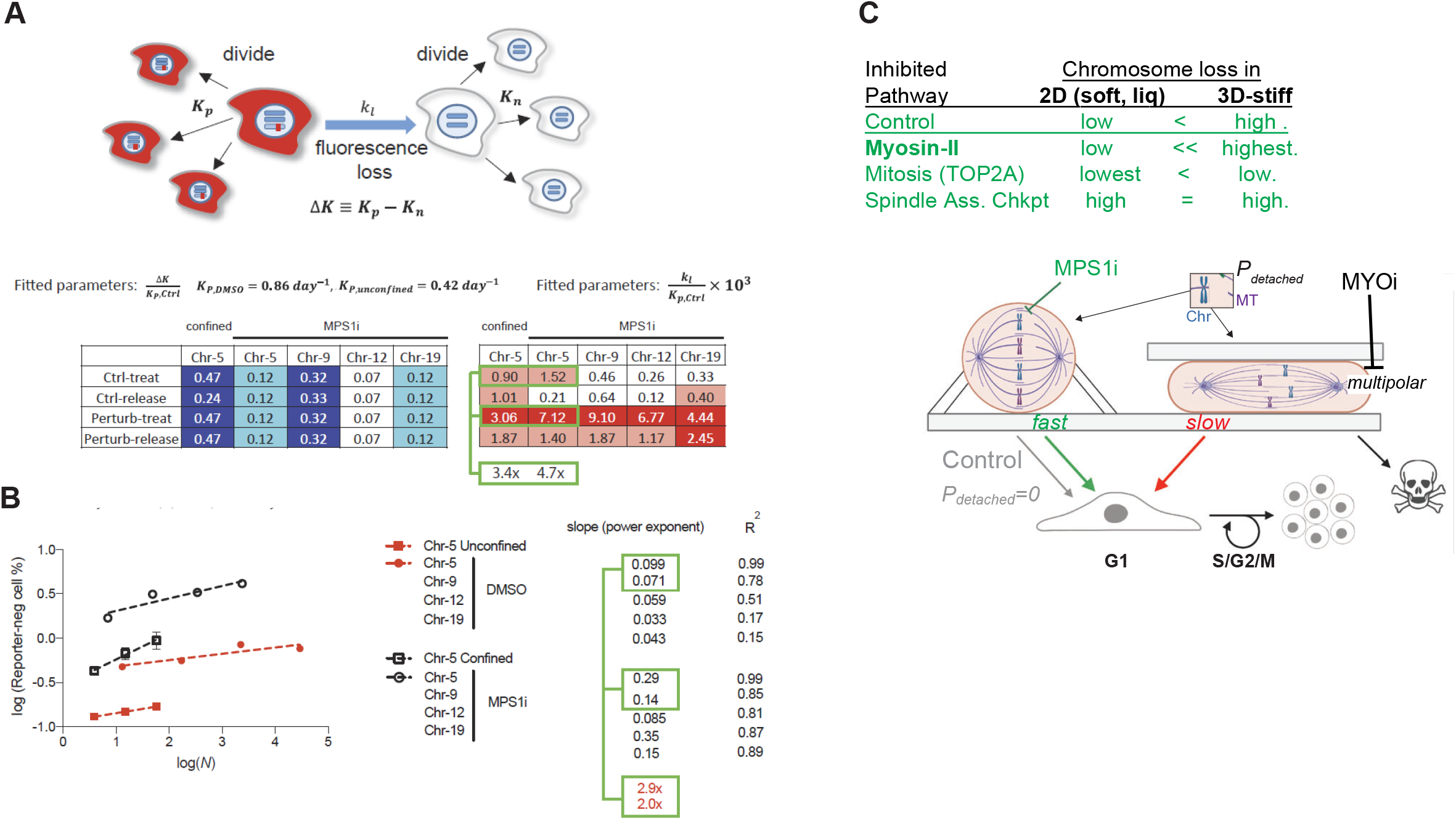
Mathematics of a ‘Heritable Loss Model’ (HLM) for ChReporter loss kinetics, including power-law scaling with cell division, and Confinement-dependent Pharmacology. **(A)** Schematic showing cell division and fluorescence loss, with *K*_*p*_, *K*_*n*_, *k*_*l*_ respectively quantifying net proliferation of reporter-pos cells, net proliferation of reporter-neg cells, and fluorescence loss rate. The latter is then derived as r(t) (see Materials and Methods). Fitting parameters are normalized to the fixed value *K*_*p,Ctrl*_ (Fig. 4G,H). We assume *ΔK* is identical for all the phases except for the MPS1i treatment phase. For each reporter *k*_*l, Perturb*_ > *k*_*l, Ctrl*_, consistent with increased Chr loss. **(B)** Power-law scaling of all ChReporters during MPS1i treatment as well as for Chr-5 reporter under 3D-rigid confinement. The perturbed processes have steeper slopes than their corresponding controls, with fold changes of 1.5 to 10. **(C)** ChReporter loss in 2D versus 3D-confinement for various pathways. Myosin-IIA suppression has no effect in 2D but increases loss in confinement. MPS1i only affects 2D, and gives results similar to rigid confinement, perhaps by limiting microtubule attachment/signaling. In all cases *some* viable, cycling cells show heritable loss.

## DISCUSSION

Myosin-II suppression amplifies confinement-induced Chr loss but despite previous speculations [20-25]. has no effect in standard 2D (**Fig.5C**). Inhibiting mitosis (with Etop) suppresses Chr loss in 2D and slightly less so in confinement, which indicates divisions *N* might be necessary but high *N* is insufficient for loss. Inhibitors of the spindle assembly checkpoint (SAC) increase ChReporter loss only in 2D culture, perhaps because in 3D-rigid confinement the added pharmacological stress favors death. Our new mono-allelic ChReporter approach to quantifying chromosome loss is more broadly applicable to cell biology and *in vivo* tests, and supports the hypothesis here that mitosis within confining microenvironments causes not just massive cell death or senescence but also *heritable mechanogenetic* changes. Stress-driven, stiffness-associated generation of rare GFP/RFP-negative cells (∼1%) of varying long-term viability can thus be directly seen *in vitro* and *in vivo*. By this method, evolutionary selection via death, stasis, or colony formation need not be inferred *per* standard genetic analyses that require killing cells and that lack key spatial information. Such analyses nonetheless support our ChReporter approach and include single cell sequencing of 100’s of cells. Similar analyses of normal unedited tissue cells also show rare losses or gains in cells within 3D microenvironments. The results of confinement here might relate – with further study – to the proposed effects of abnormal tissue architecture *via* integrins [50]. Interphase mechanosensing depends often on actomyosin connections to integrin adhesions [1-8], and the fact that myosin-II suppression limits centriole splitting [20] provides a key mechanistic explanation for why suppression in mitosis rescues “lethal” multipolars toward more viable mis-segregations (**Fig.2D,5C**).

Viable genetic changes are quickly visible here after confined mitosis, with 3D compression known to distort the MT spindle and increase Chr mis-segregation [17, 18, 46] – but the lack of effect of ‘tumor suppressor’ Myosin-II in standard 2D cultures differs from prior speculations based on 2D cultures [21-25]. The Myosin-II dependent centriole splitting mechanism [20] is nonetheless relevant to confinement effects when also considering that multipolars are “lethal” [18]. Conversely, the SAC clearly limits Chr loss in 2D but seems unimportant to Chr loss in 3D confinement. Various SAC genes such as *MPS1* are not widely identified as tumor suppressors, and this includes a recent in-depth analysis of 21 solid and liquid tumor types that concluded whole-chromosome loss is ‘driven’ by loss of tumor suppressors such as “guardian of the genome” p53 (on Chr-17) [15]. SAC genes could be essential but are also not identified in studies showing 3D-spheroids are better than 2D cultures in revealing growth effects of tumor suppressors and oncogenes [51]; in contrast, myosin-IIA (*MYH9*) was categorized as a ‘3D’ tumor suppressor. Myosin-IIA might directly regulate p53 and cause the presumed genetic changes that lead to cancer in Myosin-IIA knockdown mouse skin, but mouse tongue results seem to differ [13, 14]. In standard 2D-cultures of a cardiomyocyte tumor line, nonmuscle myosin-II depletion increased abnormal mitosis and cell death as well as MT acetylation-stabilization [52]; such a process has been linked to a phosphatase that inhibits both contractility and a MT-targeting de-acetylase [53] but also – intriguingly – to aneuploidy in breast cancer [54]. Although Chr loss here with Myosin-IIA depletion in 3D-confinement seems distinct from MT processes and 2D processes (including SAC regulation), our novel ChReporter approach might allow molecular mechanisms to be further addressed by specifically interrogating viable cells with clear and heritable Chr losses.

## SUPPLEMENT

Figure S1 shows Purification of ChReporter-positive cells and Validation of ChReporter-negatives for different chromosomes across numerous cell lines. Figure S2 shows Confined mitosis compacts mitotic chromatin, suppresses growth, and increases death. Figure S3 shows Pharmacology and confined mitosis: inhibitors such as MPS1i of Spindle Assembly Checkpoint (SAC) exert similar but distinct effects in terms of death, kinetics, and ChReporter loss.

## Materials and Methods

### Cell lines and culture

The following cancer cell lines for this study were: A549 lung adenocarcinoma, U2OS osteosarcoma, and NCI-H23 lung adenocarcinoma (referred to as H23 in text). The A549 and U2OS cell lines were obtained from the American Type Culture Collection (ATCC). The H23 cell line was a kind gift from Dr. Michael C. Bassik (Stanford University). A549 cells were cultured in Ham’s F-12 media (Gibco 11765047); U2OS cells in DMEM (Gibco, Catalog no. 10569010); and H23 cells in RPMI 1640 (Gibco, Catalog no. 11879020). The original A549 RFP-LMNB1 cell line was engineered by Sigma-Aldrich. HEK293T cells used, acquired from ATCC, for lentiviral packaging were also cultured in DMEM. All aforementioned cell lines were cultured in media supplemented with 10% (v/v) fetal bovine serum (FBS; MilliporeSigma, Catalog no. F2442) and 100 U ml^−1^ penicillin-streptomycin (Gibco, Catalog no. 15140122). All cells were passaged every 2-3 days using 0.05% Trypsin/EDTA (Gibco, Catalog no. 25300054). All cell lines were incubated at 37°C and maintained at 5% CO_2_.

The following induced pluripotent stem cell (iPSC) lines were also used, all of which were acquired from the Coriell Institute for Biomedical Research and generated/validated by the Allen Institute for Cell Science: iPSC GFP-LMNB1 (AICS-0013 cl.210), iPSC RFP-LMNB1 GFP-SEC61B (AICS-0059 cl.36), and iPSC FBL-GFP NPM1-RFP (AICS-0084 cl.18). iPSCs were cultured in mTseR Plus medium (STEMCELL Technologies, Catalog no. 05825), with mTser Plus 5X supplement and 100 U ml^−1^ penicillin-streptomycin. For passaging and maintenance of iPSCs, cells were lifted with accutase (Sigma, Catalog no. A6964) at 37°C and re-plated into 10-cm plates (Corning) coated with Matrigel (Corning, Catalog no. 356231) following the Allen Institute of Cell Science’s protocol. 10 mM ROCK inhibitor (Y-27632; STEMCELL Technologies, Catalog no. 72302) was added to replated cultures to help with adherence and to prevent differentiation. Passaging was done once iPSC cultures reached 70% confluency to prevent spontaneous differentiation. All iPSC lines were also cultured at 37°C and maintained at 5% CO_2_.

### Monoallelic chromosome tagging

All knock-in reporter lines were generated following the protocol established in (1) using CRISPR/Cas9 technology. Donor plasmids were designed such that unique designs for each target locus contain 5’ and 3’ homology arms (1 kb each) for the desired insertion site, based on the GRCh38 reference human genome. For all attempted monoallelic chromosome reporters as described in Fig. 1E, donor constructs were: AICSDP-8:TOMM20-mEGFP (Addgene plasmid #87423; http://n2t.net/addgene:87423; RRID:Addgene_87423), AICSDP-13:FBL-mEGFP (Addgene plasmid #87427; http://n2t.net/addgene:87427; RRID:Addgene_87427), AICSDP-35:AAVS1-mEGFP (Addgene plasmid #91565; http://n2t.net/addgene:91565; RRID:Addgene_91565), AICSDP-42:AAVS1-mTagRFPT-CAAX (Addgene plasmid #107580; http://n2t.net/addgene:107580; RRID:Addgene_107580), AICSDP-1:PXN-EGFP (Addgene plasmid #87420; http://n2t.net/addgene:87420; RRID:Addgene_87420), AICSDP-10:LMNB1-mEGFP (Addgene plasmid #87422; http://n2t.net/addgene:87422; RRID:Addgene_87422), AICSDP-52: HIST1H2BJ-mEGFP (Addgene plasmid #109121; http://n2t.net/addgene:109121 ; RRID:Addgene_109121), AICSDP-7:SEC61B-mEGFP (Addgene plasmid # 87426; http://n2t.net/addgene:87426; RRID:Addgene_87426).

For editing, we use the ribonucleic protein (RNP) method with recombinant wild type *S. pyogenes* Cas9 protein pre-complexed with a synthetic CRISPR RNA (crRNA) and a trans-activating crRNA (tracrRNA) duplex. Recombinant wild-type Cas9 protein was purchased from the University of California–Berkeley QB3 Macrolab, while crRNA and tracrRNA oligonucleotides were designed by and purchased from Horizon Discovery. For transfection of donor templates into target cells, we used the electroporation using a Gene Pulser Xcell Electroporation System (Bio-Rad). 700,000 targets cells were lifted using 0.05% Trypsin/EDTA, resuspended in 200 ul of fresh media without penicillin-streptomycin, and loaded into a 0.4-cm cuvette. 4 µL of both 10 µM crRNA:tracrRNA duplex and 10 µM recombinant Cas9 protein were added to the cell solution, as well as 8 µg of donor plasmid. Electroporation conditions were as follows: (1) A549 and H23: 200V with 45 ms pulse length using a square-wave protocol; (2) U2OS: 160V with 30 ms pulse length using a square-wave protocol. After

